# Expansion of the prime editing modality with Cas9 from *Francisella novicida*

**DOI:** 10.1101/2021.05.25.445577

**Authors:** Yeounsun Oh, Wi-jae Lee, Hanseop Kim, Lee Wha Gwon, Young-Hyun Kim, Young-Ho Park, Chan Hyoung Kim, Kyung-Seob Lim, Bong-Seok Song, Jae-Won Huh, Sun-Uk Kim, Bong-Hyun Jun, Cheulhee Jung, Seung Hwan Lee

**Affiliations:** Futuristic Animal Resource & Research Center (FARRC), Korea Research Institute of Bioscience and Biotechnology (KRIBB), Cheongju, Korea; National Primate Research Center (NPRC), Korea Research Institute of Bioscience and Biotechnology (KRIBB), Cheongju, Korea; Department of Biotechnology, College of Life Sciences and Biotechnology, Korea University, Seoul, Republic of Korea; Department of Biomolecular Science, KRIBB School of Bioscience, Korea University of Science and Technology, Gajeong-dong, Yuseong-gu, Daejeon, Republic of Korea; Department of Functional Genomics, KRIBB School of Bioscience, Korea University of Science and Technology (UST), Daejeon, Korea; School of Life Sciences and Biotechnology, BK21 Plus KNU Creative BioResearch Group, Kyungpook National University, Daegu, Republic of Korea; Department of Bioscience and Biotechnology, Konkuk University, Seoul, Korea; Department of Biological Sciences, Chungnam National University, Daejeon, Korea

## Abstract

Prime editing can induce a desired base substitution, insertion, or deletion in a target gene using reverse transcriptase (RT) after nick formation by CRISPR nickase. In this study, we developed a technology that can be used to insert or replace external bases in the target DNA sequence by linking reverse transcriptase to the *Francisella novicida* Cas 9 [FnCas9(H969A)] nickase module, which is a CRISPR-Cas9 ortholog. Using FnCas9(H969A) nickase, the targeting limitation of existing *Streptococcus pyogenes* Cas9 nickase [SpCas9(H840A)]-based prime editing was dramatically extended, and accurate prime editing was induced specifically for the target genes.

---

The recent development of reverse transcriptase (RT)-based target DNA editing technology (i.e., prime editing) is based on the SpCas9(H840A) module^1-3^. Since prime editing technology can induce various types of mutations^2,4-7^ compared to base editing^8,9^ that only induces deamination-mediated base substitutions (A to G or C to T), more than 90% of the pathogenic mutations reported to date may have been corrected without double-stranded DNA cleavage^10^. However, since SpCas9 module-based prime editing technology can insert, delete, and replace bases at a distance of 3 bp from the protospacer adjacent motif (PAM) sequence (NGG) on the target DNA, there is a PAM limitation.^11,12^ Therefore, there is still room for improvement in prime editing technology. In this study, we improved the safety and effectiveness limitations of SpCas9(H840A)-RT with a new approach using *F. novicida* Cas9^13-16^, a CRISPR-Cas9 ortholog-based RT [FnCas9(H969A)-RT], in human-derived cell lines (HEK293FT and HeLa cells).

Compared to SpCas9(H840A)^17^, FnCas9(H969A) forms a nick in the non-target strand from the PAM (NGG) but in the same target sequence [i.e., 6 bp upstream from the PAM for FnCas9(H969A) and 3 bp upstream from the PAM for SpCas9(H840A)]^14^ **(Supplementary Figure 1a)**. Therefore, the FnCas9(H969A) module has the advantage of expanding the region recognized as the reverse transcription template (RTT) following the primer binding site (PBS) sequence. We further tried to expand the target sequences **(Supplementary Figure 1b-d)** that can be used to cover each gene in human-derived cell lines using the FnCas9(H969A)-RT system **(Figure 1a)**.

**Figure 1.**
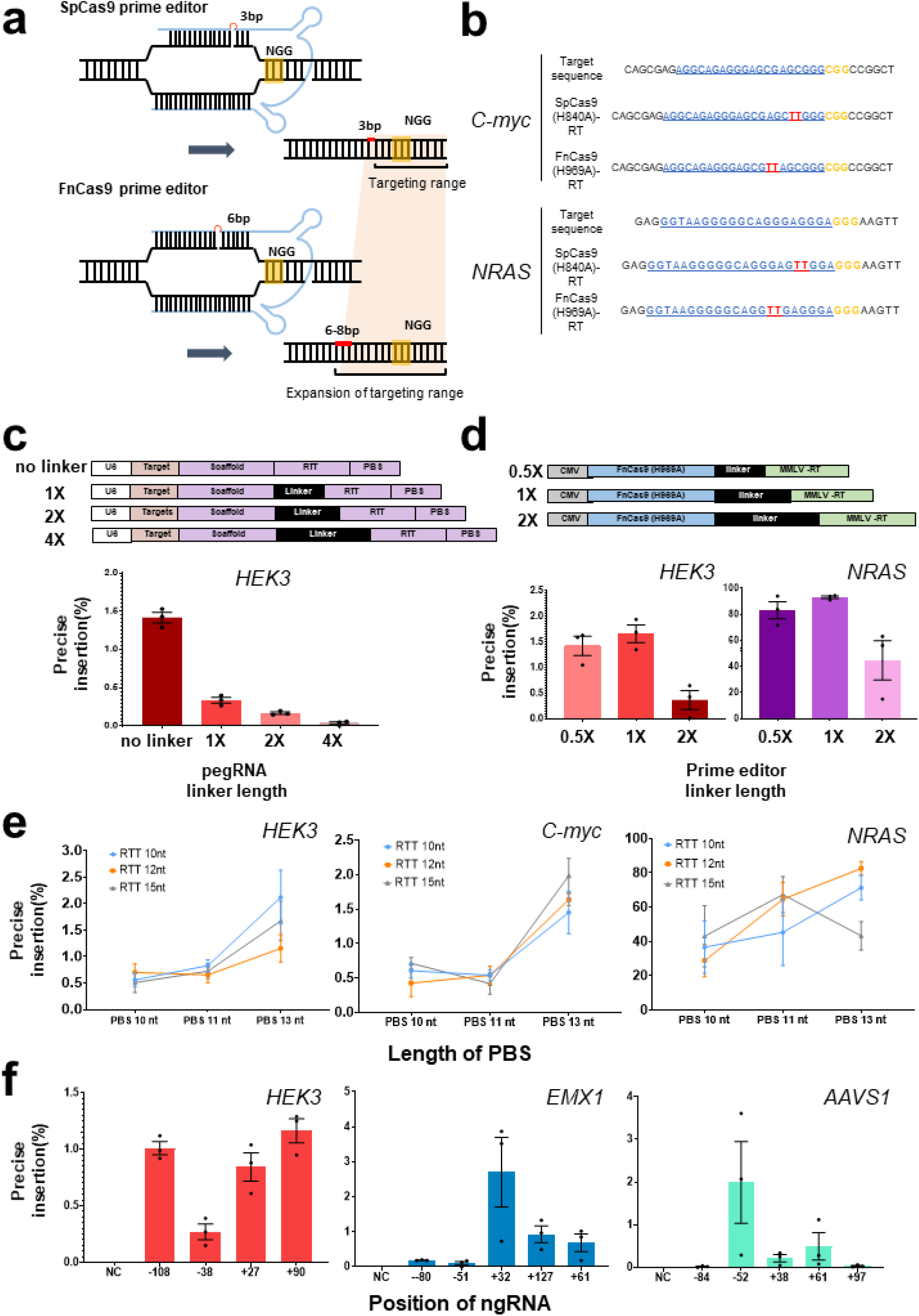
Targeted prime editing and optimization with FnCas9(H969A)-RT. **a**. Comparison between SpCas9(H840A) and FnCas9(H969A) nickase-based prime editing. SpCas9(H840A) forms a nick 3–4 bp upstream from the PAM (NGG) and FnCas9 (H969A) forms a nick 6–8 bp upstream from the PAM (NGG). Only pegRNAs are shown in blue for clarity, each PAM (NGG) sequence is shown in yellow, and the targeted base insertion is shown in red. **b**. Comparison of target-specific nucleotide insertion into the *HEK3* and *NRAS* genes according to the different nicking positions of SpCas9(H840A)-RT and FnCas9(H969A)-RT. PAM (NGG) sequences are shown in yellow, protospacers are shown in blue, and inserted base sequences are shown in red. **c**. Preparation of various pegRNAs for optimizing the prime editing efficiency and comparison of nucleotide insertion efficiency according to linker length (no linker, 1×, 2×, 4× linkers) for *HEK3*. **d**. Construction of FnCas9(H969A)-RT with linkers of various lengths (0.5×, 1×, 2×) and comparison of nucleotide insertion efficiency for *HEK3* and *NRAS*. **e**. Comparison of gene editing efficiency of FnCas9(H969A)-RT according to the PBS and RTT length in pegRNA on various genes (*HEK3, C-myc, NRAS*). **f**. Comparison of base insertion efficiency according to ngRNA targeting at various positions in the PE3 method of FnCas9 (H969A)-RT. Each histogram was plotted by applying standard error of the mean values to repeated experimental values (n = 3). PBS: primer binding site; RTT: reverse transcription template; pegRNA: prime editing guideRNA; MMLV-RT: reverse transcriptase domain from Moloney Murine Leukemia Virus.

First, an RT enzyme was connected to the FnCas9(H969A) nickase module to optimize the performance of the prime editor on the target gene. To optimize the efficiency of the newly applied FnCas9 prime editor, FnCas9(H969A)-RT or prime editing guide RNA (pegRNA) with various linker lengths were prepared **(Figure 1b,c top, Supplementary Table 1)**. The pegRNA was constructed to allow for insertion of a “TT” di-nucleotide sequence at the expected cleavage point [6 and 3 nucleotides upstream from the PAM (NGG) sequence for Fn and Sp, respectively] for the *HEK3* site by considering the nicking point on each target sequence based on SpCas9 or FnCas9 nickase modules. The cell lines were then co-transfected with pegRNA/FnCas9(H969A)-RT and nicking-guide RNA (ngRNA) expression plasmids **(Supplementary Figure 1e)**. This confirmed that the bi-nucleotide TT was accurately inserted in the genomic DNA (*C-myc, NRAS*) at a site 6 nucleotides away from the PAM in the target sequence **(Figure 1b)**. The pegRNA without a linker showed the highest efficiency in *HEK3* locus **(Figure 1c)**, and optimized TT insertion was achieved when using the FnCas9 prime editor with a 1× linker length for *HEK3* and *NRAS* locus **(Figure 1d)**.

To confirm the effect of the length of the PBS and RTT in pegRNA on prime editing efficiency, several candidate pegRNAs with different lengths of PBS and RTT were designed and produced **(Supplementary Table 1)**, and a TT insertion was attempted at various genes (*HEK3, C-myc, NRAS*) in human cell lines **(Figure 1e, Supplementary Figure 1f, h, j)**. The external TT insertion efficiency at the *HEK3* site increased slightly as the length of the PBS increased **(Figure 1e, Supplementary Figure 1g, i, k)**. FnCas9(H969A)-RT resulted in 0.99% (*HEK3*), 0.92% (*C-myc*), and 53.7% (*NRAS*) correct TT insertion [6 bp upstream from PAM (NGG)] compared to SpCas9(H840A)-RT on average **(Figure 1e, Supplementary Figure 1g, i, k)**, and the efficiency varied according to the length of RTT. Notably, the TT insertion efficiency was significantly different for each targeted gene (*HEK3, EMX1, AAVS1*) according to the location of the nicking guide RNA **(Figure 1f, Supplementary Table 1)** on the target-strand side when applying the prime editing with target-strand nicking (PE3) approach **(Supplementary Figure 1e)**. To directly compare the efficiency of FnCas9(H969A)-RT with that of SpCas9(H840A)-RT for various genes, another TT insertion experiment was performed for the same target sequence in various genes (*C-myc, NRAS*) with both prime editing systems **(Supplementary Figure 2)**. Next-generation sequencing **(Supplementary Table 3)** showed that compared to SpCas9-RT (29.34-95.18%), FnCas9-RT exhibited greater variation in TT insertion efficiency (3.82-87.56%), which varied depending on the target gene **(Supplementary Figure 2a)**. Although TT insertion was accurately induced for a given target sequence, unintended indel formation caused by the PE3 correction method occurred at a frequency of 4.79% (*NRAS*) and 1.13% (*C-myc*) when using SpCas9(H840A)-RT, and at a frequency of 1.32% (*NRAS*) and 1.84% (*C-myc*) in the case of FnCas9(H969A)-RT. These results may reflect the weaker nicking property of FnCas9(H969A) compared to SpCas9(H840A) **(Supplementary Figure 2d-f)**. Indeed, when analyzing the off-target effect of the FnCas9-based prime editor for each target gene, we found that no significant off-target insertion was observed by FnCas9(H969A)-RT **(Supplementary Figure 2b, Supplementary Table 4)**. Moreover, although FnCas9(H969A)-RT could induce TT insertion by prime editing in the HeLa cell line, the overall targeted insertion efficiency was low and was more sensitive than that of SpCas9(H840A)-RT **(Supplementary Figure 2c)**.

Because the FnCas9 nickase (H969A) module induces nicks on the non-target strand side of the target gene in contrast to the SpCas9 nickase (H840A) module **(Supplementary Figure 1a)**, reverse transcription can be initiated by RT from a position farther away from the nucleotide sequence recognized as the PAM (NGG). According to this difference, we directly compared the range that can induce prime editing based on FnCas9-RT and SpCas9-RT with respect to the same target sequence **(Figure 2a–c)**. In HEK293FT cells, the efficiency of prime editing was first compared for the same target sequence in the *C-myc* and *EMX1* genes, and the point at which the base is inserted (AA for *C-myc*, TT for *EMX1*) was indicated according to the point at which nicks were formed (+1) by the SpCas9 nickase (H840A) module **(Figure 2a)**. SpCas9 nickase(H840A)-RT showed a correction efficiency of 27–39% for the AA insertion at *C-myc* from the –1 to +3 position in the sequence based on the nick position **(Figure 2b,c, top)**. By contrast, FnCas9 nickase(H969A)-RT showed a 2–8% insertion efficiency from position −6 to +3 based on the nick position **(Figure 2b,c, bottom)**. This result showed that based on the shared PAM sequence (NGG) in the same target, FnCas9(H969A)-RT greatly expanded the prime editing range. In particular, FnCas9(H969A)-RT can effectively induce prime editing of various point mutations or insertions in regions that cannot be edited using the SpCas9 system **(Figure 2d, Supplementary Figure 2g)**. Similar results were found for the TT insertion efficiency for the same target sequence in the *EMX1* gene **(Supplementary Figure 3)**, in which the range of genome editing by FnCas9(H969A)-RT was significantly expanded compared to that of SpCas9(H840A)-RT editing.

**Figure 2.**
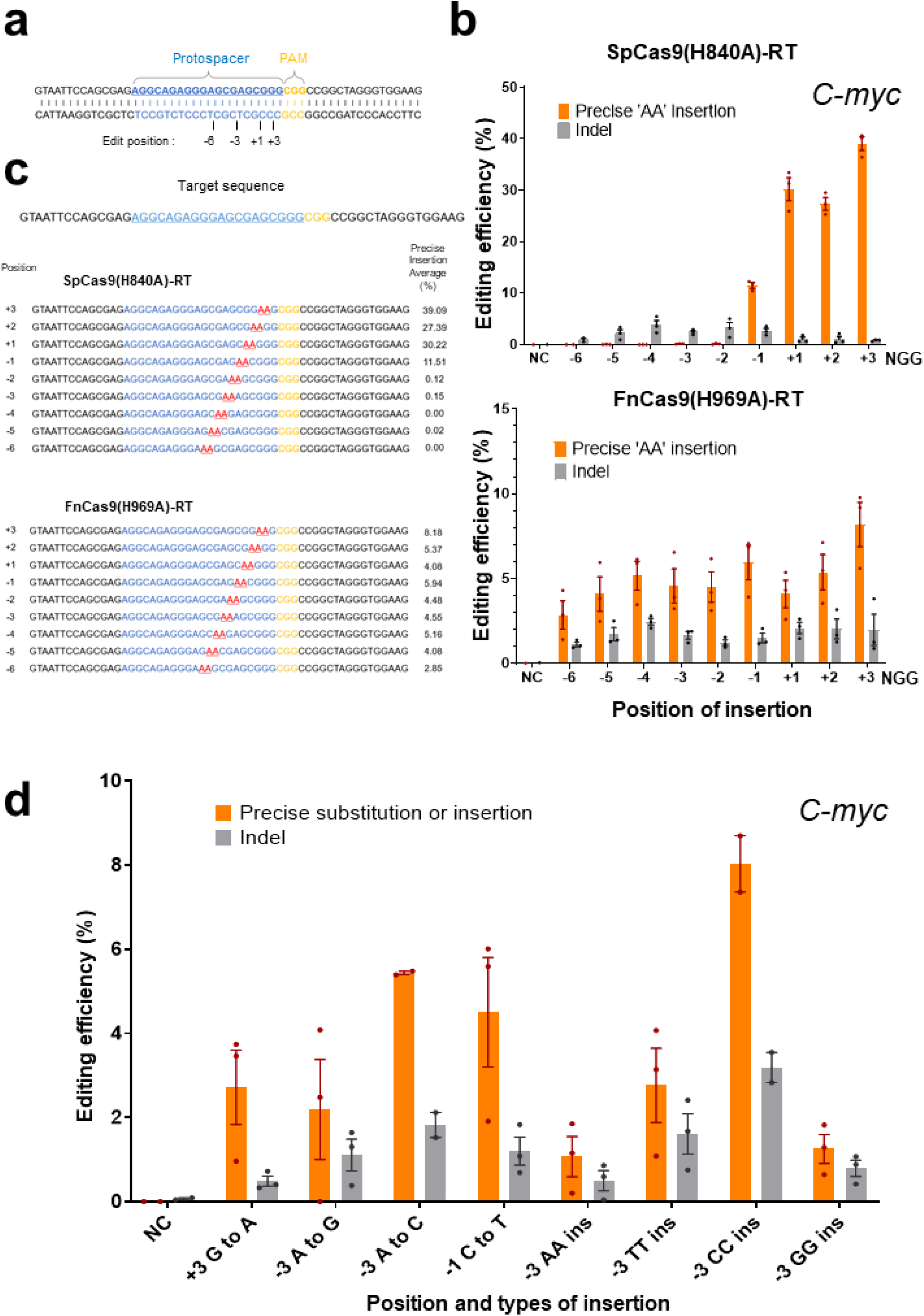
Expansion of the range of prime editing by FnCas9(H969A)-RT. **a**. Schematics of the protospacer (blue), PAM (yellow), and base insertion site (indicated numbers below the target sequence) on the target gene (*C-myc*) recognized by SpCas9(H840A)-RT and FnCas9(H969A)-RT. **b**. Direct comparison of the efficiency of base insertion by SpCas9(H840A)-RT and FnCas9(H969A)-RT. Top: Comparison of efficiency according to the base insertion site caused by SpCas9(H840A)-RT. Bottom: Comparison of efficiency according to the base insertion site caused by FnCas9(H969A)-RT. **c**. Next-generation sequencing result of site-specific base insertion induced by SpCas9(H840A)-RT and FnCas9(H969A)-RT. Protospacers are shown in blue, PAM (NGG) sequences are shown in yellow, and inserted sequences (AA) are shown in red. Each position and efficiency in which the AA bases are inserted are indicated to the left and right of the target sequence, respectively. **d**. Prime editing in various positions and types caused by FnCas9(H969A)-RT. The editing efficiency according to each position and type of prime editing is plotted in the histogram.

Finally, we examined whether the optimized FnCas9(H969A)-RT can be used in combination with SpCas9(H840A)-RT to induce prime editing simultaneously on the target gene **(Figure 3)**. The TT insertion was respectively induced by FnCas9(H969A)-RT and SpCas9(H840A)-RT in two adjacent nucleotide sequences on the *NRAS* gene **(Figure 3b)**. To increase the efficiency of simultaneous editing of the two sites, nickases on the target-strand side were treated at various positions, resulting in different editing efficiencies **(Figure 3a)**. In samples treated with both prime editors simultaneously, the TT insertion was generally induced with high efficiency (25.81–47.06%) by SpCas9(H840A)-RT, whereas that induced by FnCas9(H969A)-RT showed more varied efficiency (0.12–44%) according to the nickase position on the target-strand side. The overall rate of simultaneous TT insertion by the two prime editors was relatively low (0.02–1.3%).

**Figure 3.**
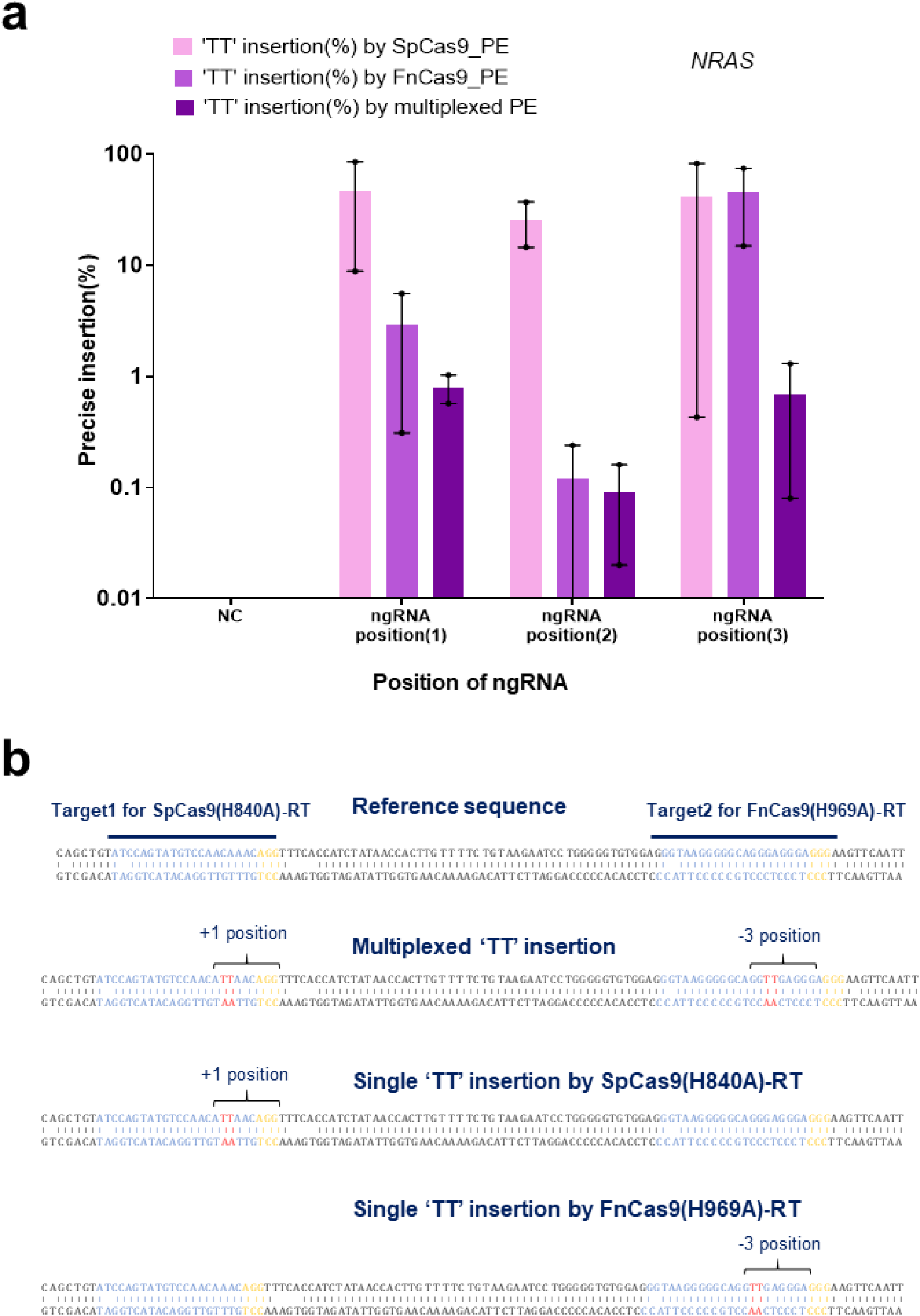
Multiplexed prime editing induced by simultaneous treatment of SpCas9(H840A)-RT and FnCas9(H969A)-RT. **a**. Comparison of base insertion efficiency of multiplexed prime editing by SpCas9(H840A)-RT and FnCas9(H969A)-RT according to ngRNA targeting at various positions (1-3, **Supplementary Figure 1d**) using the PE3 method. Each histogram was plotted by applying standard error of the mean values to repeated experimental values (n = 3). **b**. Next-generation sequencing results of multiplexed TT insertion using SpCas9(H840A)-RT and FnCas9(H969A)-RT. Each target sequence [protospacer shown in blue and PAM (NGG) shown in yellow] for SpCas9(H840A)-RT and FnCas9(H969A)-RT is indicated. The targeted TT insertion is shown in red and each position is shown above the base. PE3: prime editing with target-strand nicking; ngRNA: nicking guide RNA.

In summary, we developed a new prime editing module that can dramatically expand the capabilities of SpCas9-based prime editing technology. Compared to the SpCas9(H840A)-RT prime editor, FnCas9(H969A)-RT, which recognizes the same PAM (NGG) nucleotide sequence, induces cleavage of the non-target strand at a further distance from the PAM, thereby expanding the range in which the RT can operate. In addition, prime editing could be achieved with both modules simultaneously. The FnCas9(H969A)-RT-based prime editing technique developed in this study can be further improved by expanding the PAM or by increasing the editing efficiency using protein engineering in the near future.

## ACKNOWLEDGMENTS

This research was supported by grants from the National Research Foundation funded by the Korean Ministry of Education, Science and Technology (NRF-2019R1C1C1006603); the Technology Innovation Program funded by the Ministry of Trade, Industry & Energy (MOTIE, Korea) (20009707); and KRIBB Research Initiative Program (KGM1051911, KGM4252122, KGM5382113, KGM4562121, KGM5282113).

## AUTHOR CONTRIBUTIONS

Conceptualization: Y.O., W.L., and S.H.L.; Methodology, Y.O., W.L., H.K., L.W.G., and S.H.L.; Software, Y.H.K., Y.H.P., and S.H.L.; Validation, Y.O. and C.H.K.; Formal Analysis, Y.O., W.L. and S.H.L.; Investigation, Y.O., W.L., K.S.L., S.B.S., J.W.H., S.U.K., and S.H.L.; Resources, Y.H.K., Y.H.P., K.S.L., and B.H.J.; Data Curation, Y.O., W.L., and S.H.L.; Writing-Original Draft, Y.O., W.L., C.J., and S.H.L.; Writing-Original Draft, Y.O., W.L., C.J., and S.H.L.; Visualization, Y.O., W.L., and S.H.L.; Supervision, C.J. and S.H.L.

## COMPETING INTERESTS

The authors declare no competing interests.

## METHODS

### Purification of SpCas9 and FnCas9 proteins

Subcloned pET28a-SpCas9 and pET28a-FnCas9 bacterial expression vectors were transformed into *Escherichia coli* BL21(DE3) cells, respectively, and colonies were grown at 37°C. After growing to an optical density of 0.6 in a 500 ml culture flask, induction was performed at 18°C for 48 h by isopropyl β-D-1-thiogalactopyranoside treatment. The cells were then precipitated by centrifugation and resuspended in lysis buffer [20 mM Tris-HCl (pH 8.0), 300 mM NaCl, 10 mM β-mercaptoethanol, 1% Triton X-100, and 1 mM phenylmethylsulfonyl fluoride(PMSF)]. The cells were disrupted by sonication on ice for 3 min, and the cell lysates were separated by centrifugation at 20,000 ×*g* for 10 min. The purification process was carried out as previously described^18^. Each purified SpCas9 and FnCas9 protein was replaced with a Centricon filter (Amicon Ultra) as a storage buffer [200 mM NaCl, 50 mM HEPES (pH 7.5), 1 mM dithiothreitol (DTT), and 40% glycerol] for long-term storage at −80°C. The purity of the purified proteins was confirmed by sodium dodecyl sulfate (SDS)-polyacrylamide gel electrophoresis (8–10%), and protein activity was tested using an *in vitro* polymerase chain reaction (PCR) amplicon cleavage assay.

### *In vitro* transcription for guide RNA synthesis

For *in vitro* transcription, DNA oligos containing an sgRNA sequence **(Supplementary Table 2)** corresponding to each target sequence were purchased from Cosmo Genetech. After extension PCR (denaturation at 98°C for 30 s, primer annealing at 62°C for 10 s, and elongation at 72°C for 10 s, 35 cycles) using the DNA oligos, DNA was purified using GENECLEAN® Turbo Kit (MP Biomedicals). The purified template DNA was mixed with an *in vitro* transcription mixture [T7 RNA polymerase (NEB), 50 mM MgCl_2_, 100 mM rNTPs (rATP, rGTP, rUTP, rCTP), 10× RNA polymerase reaction buffer, murine RNase inhibitor (NEB),100 mM DTT, and DEPC) and incubated at 37°C for 8 h. The DNA template was completely removed by incubation with DNase I (NEB) at 37°C for 1 h, and the RNA was further purified using GENECLEAN® Turbo Kit (MP Biomedicals) for later use. The purified RNA was concentrated by lyophilization (20,000 ×*g* at −55 °C for 1 h) and stored at −80°C.

### *In vitro* cleavage assay and DNA sequencing for nicking point analysis

Using the purified FnCas9 or SpCas9 recombinant protein, an *in vitro* cleavage assay was performed to determine the location of nicks in the non-target strand of the DNA sequence to be prime-edited. Each target site was cloned into a T-vector, and then T-vectors were purified and incubated in cleavage buffer (NEB3, 10 μl) with FnCas9 at 37°C for 1 h. The cleavage reaction was stopped by adding a stop buffer (100 mM Ethylenediaminetetraacetic (EDTA) acid, 1.2% SDS), and only the plasmid DNA was separated through the column (Qiagen, QIAquick^®^ PCR purification Kit).

The cleavage point of FnCas9 or SpCas9 was confirmed by run-off sequencing analysis of the cut fragments. The last nucleotide sequence was confirmed by the A-tailing of polymerase and the cleavage pattern was analyzed by comparative analysis with reference sequences.

### Design and cloning of the prime editor and pegRNA expression vectors

The cytomegalovirus promoter-based SpCas9(H840A)-RT and FnCas9(H969A)-RT expression vectors were constructed to induce genome editing in human cell lines. To optimize the efficiency of the newly developed FnCas9 prime editor, FnCas9 prime editors with various linker lengths were prepared or the linker length of pegRNA was diversified **(Supplementary Table 1)**. In addition, pegRNA containing the corresponding nucleotide sequence (TT or AA) was prepared so that the nucleotide could be inserted into the target nucleotide sequence (*HEK3, AAVS1, C-myc, NRAS, EMX1*). The PBS and RTT regions in pegRNA were designed and produced according to the non-target strand nick site generated by the FnCas9 nickase module **(Supplementary Figure 1a)**. As a positive control, the pegRNA of SpCas9-RT was constructed so that a TT base was inserted 3 bp in front of the PAM in consideration of the nicking point. By contrast, in FnCas9-RT, pegRNA was constructed so that the TT base was inserted at the expected cleavage point (6 bp upstream from the PAM). The pegRNA expression vector is driven by the *U6* promoter, and was designed and manufactured so that only the protospacer, PBS, and RTT can be replaced with restriction enzymes according to the target nucleotide sequence.

### Cell culture and transfection

Human-derived cell lines (HEK293FT and HeLa) were purchased from Invitrogen (R70007) and American Type Culture Collection (CCL-2), respectively. The cells were maintained in Dulbecco’s modified Eagle’s medium (DMEM) with 10% fetal bovine serum (both from Gibco) at 37°C in the presence of 5% CO_2_. Cells were subcultured every 48 h to maintain 70% confluency. For target sequence editing, 2 × 10^5^ HEK293FT or HeLa cells were transfected with plasmids expressing pegRNAs (240 pmol), the prime editor expression plasmid [SpCas9(H840A)-RT (Addgene no. 132775), FnCas9(H969A)-RT (developed in this study)], and nicking sgRNA (60 pmol) via electroporation using an Amaxa electroporation kit (V4XC-2032; program: CM-130 for HEK293FT cells, CN-114 for HeLa cells). In parallel with the SpCas9 prime editor, the FnCas9 prime editor was transfected targeting the same nucleotide sequence in HEK293FT cells. Transfected cells were transferred to a 24-well plate containing DMEM (500 μl/well), pre-incubated at 37°C in the presence of 5% CO_2_ for 30 min, and incubated under the same conditions for subculture.

### Purification of genomic DNA and construction of target site amplicons

Genomic DNA (gDNA) was extracted from the cultured cells 72 h after genome editing. The gDNA was isolated using DNeasy Blood and Tissue Kit (Qiagen). Target amplicons were obtained through PCR (denaturation at 98°C for 30 s, primer annealing at 58°C for 30 s, elongation at 72°C for 30 s, 35 cycles**; Supplementary Table 3**) from gDNA extracted from cells treated with each SpCas9 or FnCas9 prime editor. Targeted amplicon next-generation sequencing was performed using nested PCR (denaturation: 98°C for 30 s, primer annealing: 58°C for 30 s, elongation: 72°C for 30 s, 35 cycles) to analyze the efficiency of base (TT or AA) insertion.

### Targeted amplicon sequencing and data analysis

To prepare the targeted amplicon library, gDNA was extracted from the cells and further amplified using DNA primers **(Supplementary Table 3)**. Nested PCR (denaturation at 98°C for 30 s, primer annealing at 62°C for 15 s, and elongation at 72°C for 15 s, 35 cycles) was performed to conjugate adapter and index sequences to the amplicons. All targeted amplicon sequencing and data analysis were performed as suggested in a previous study^18^.

---

**Supplementary Figure 1.**
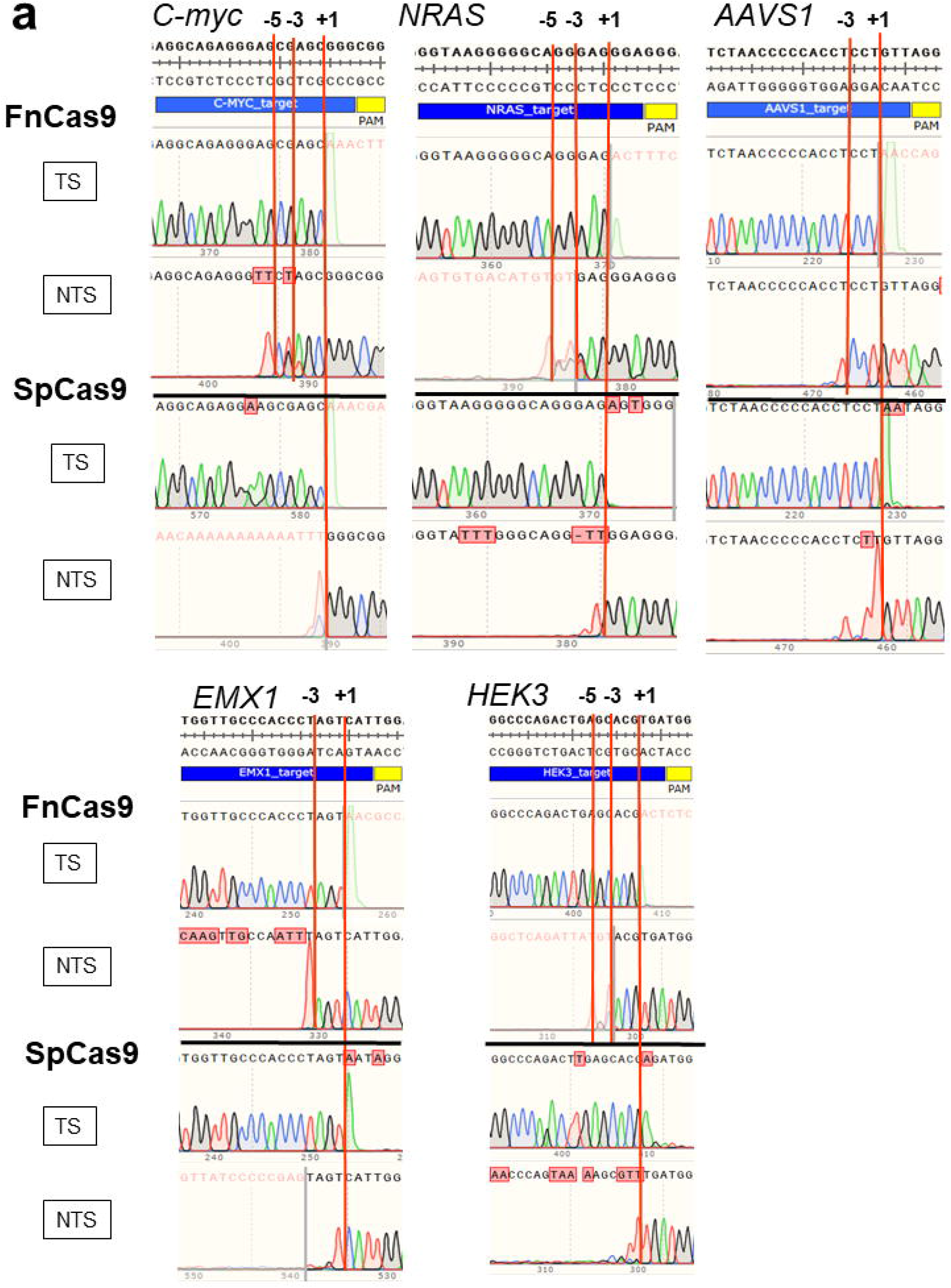
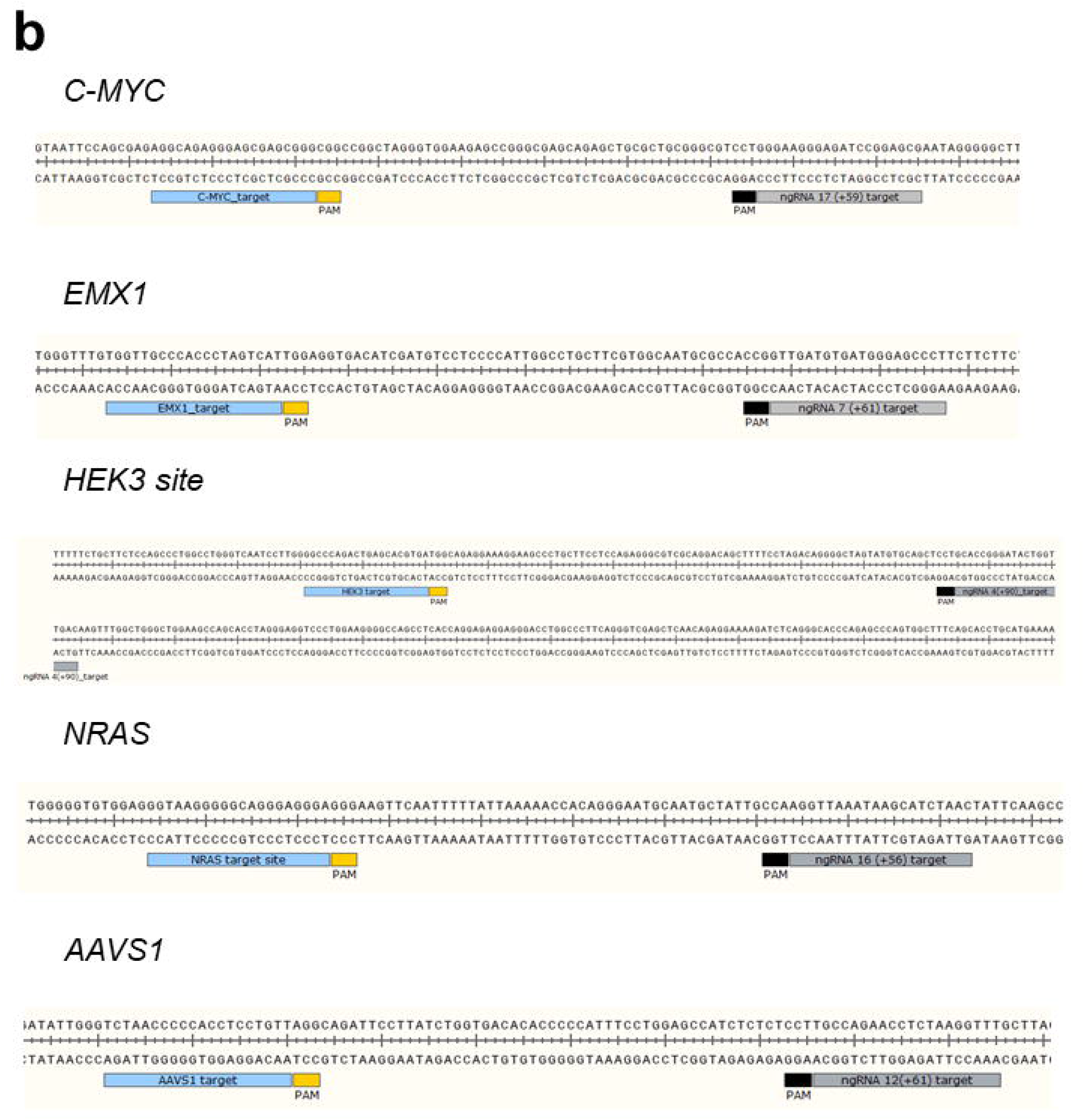
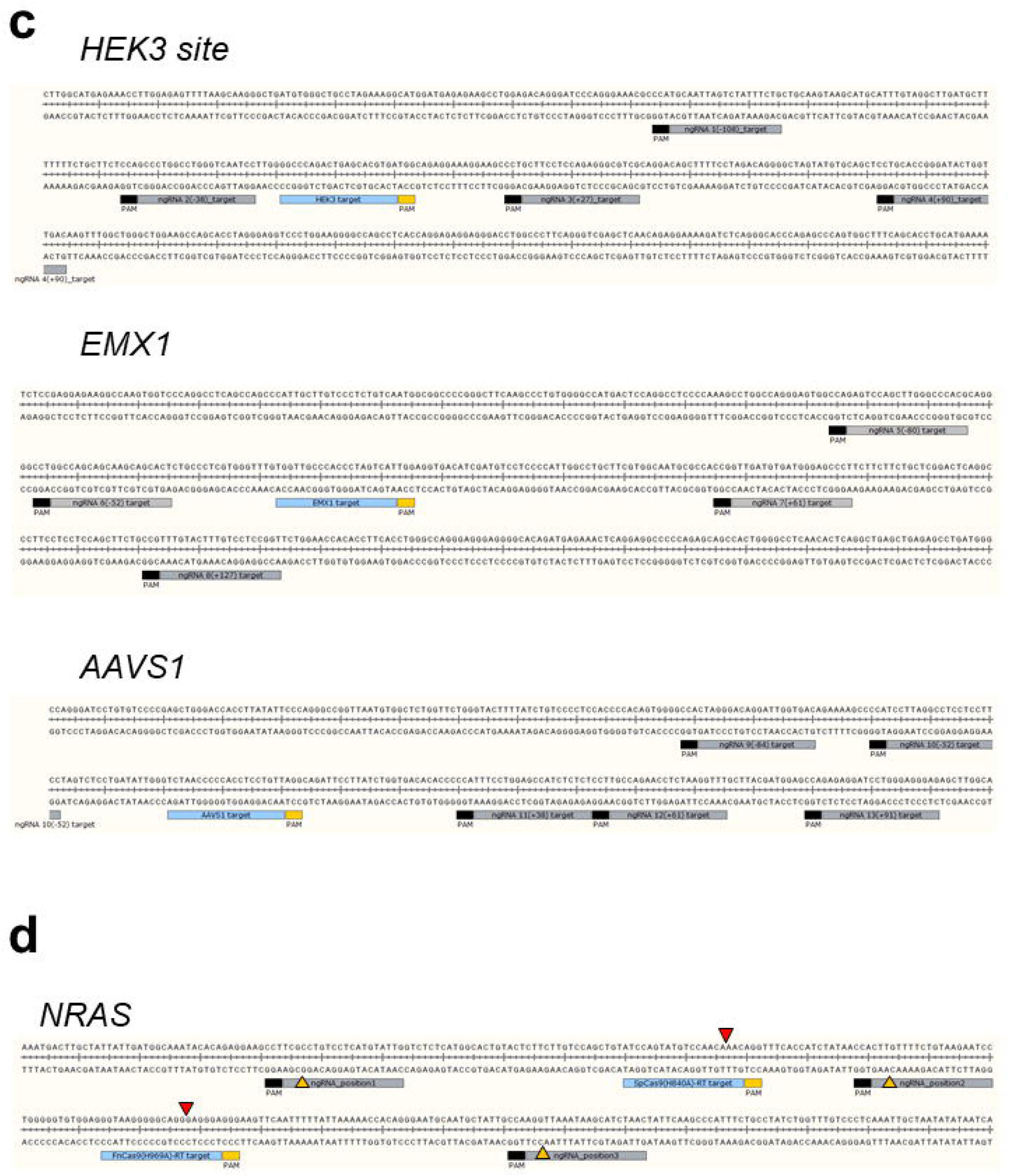
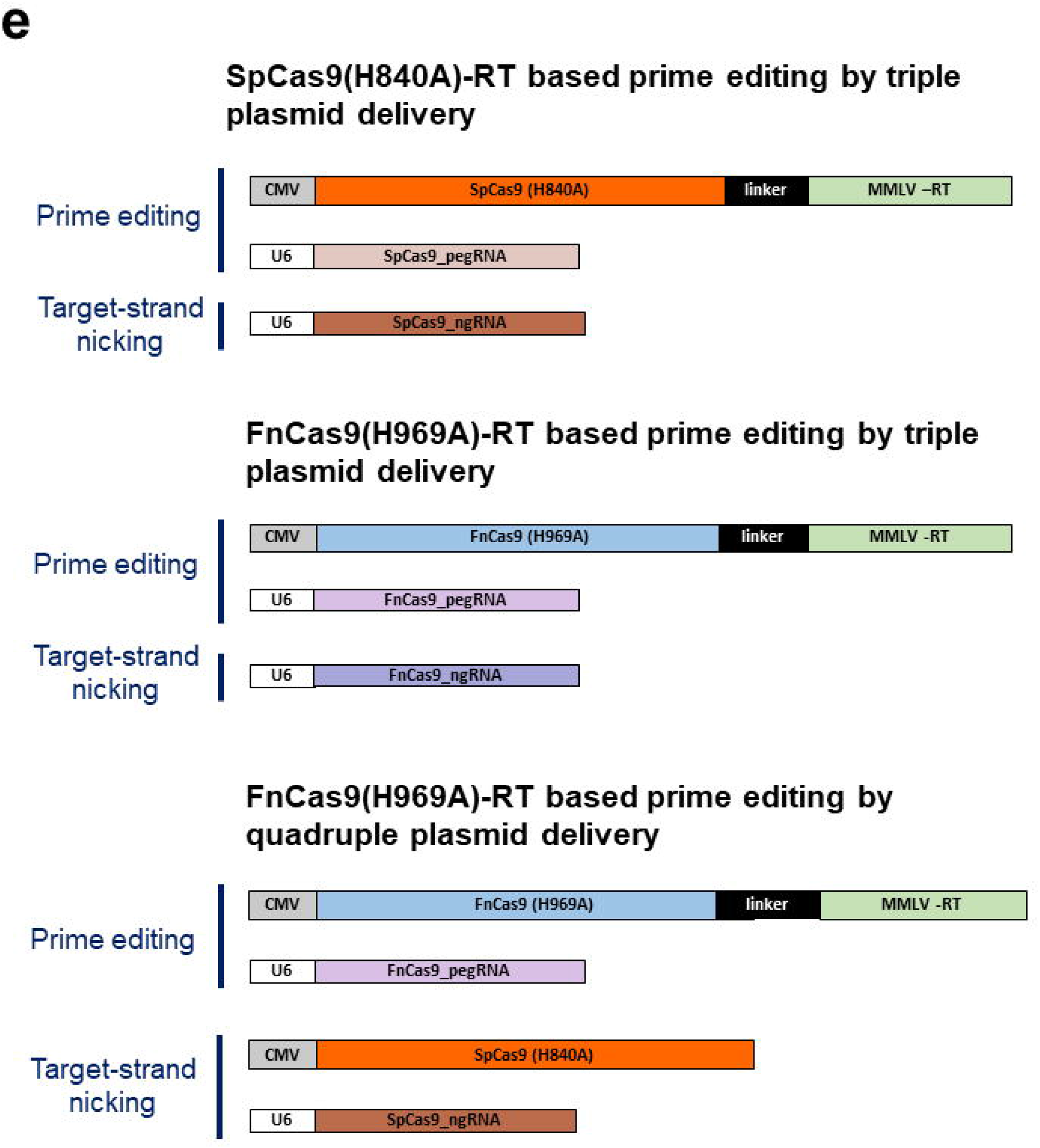
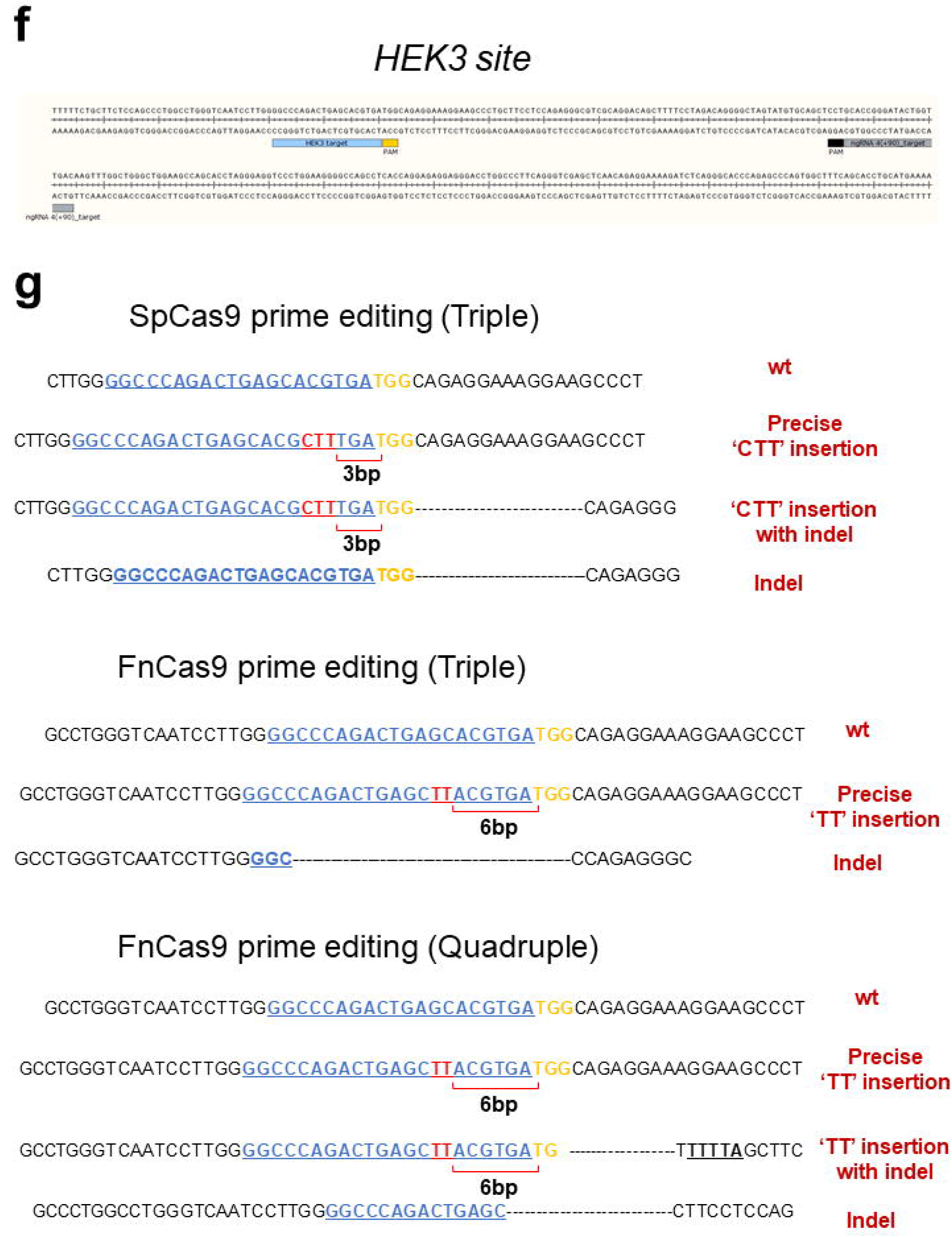
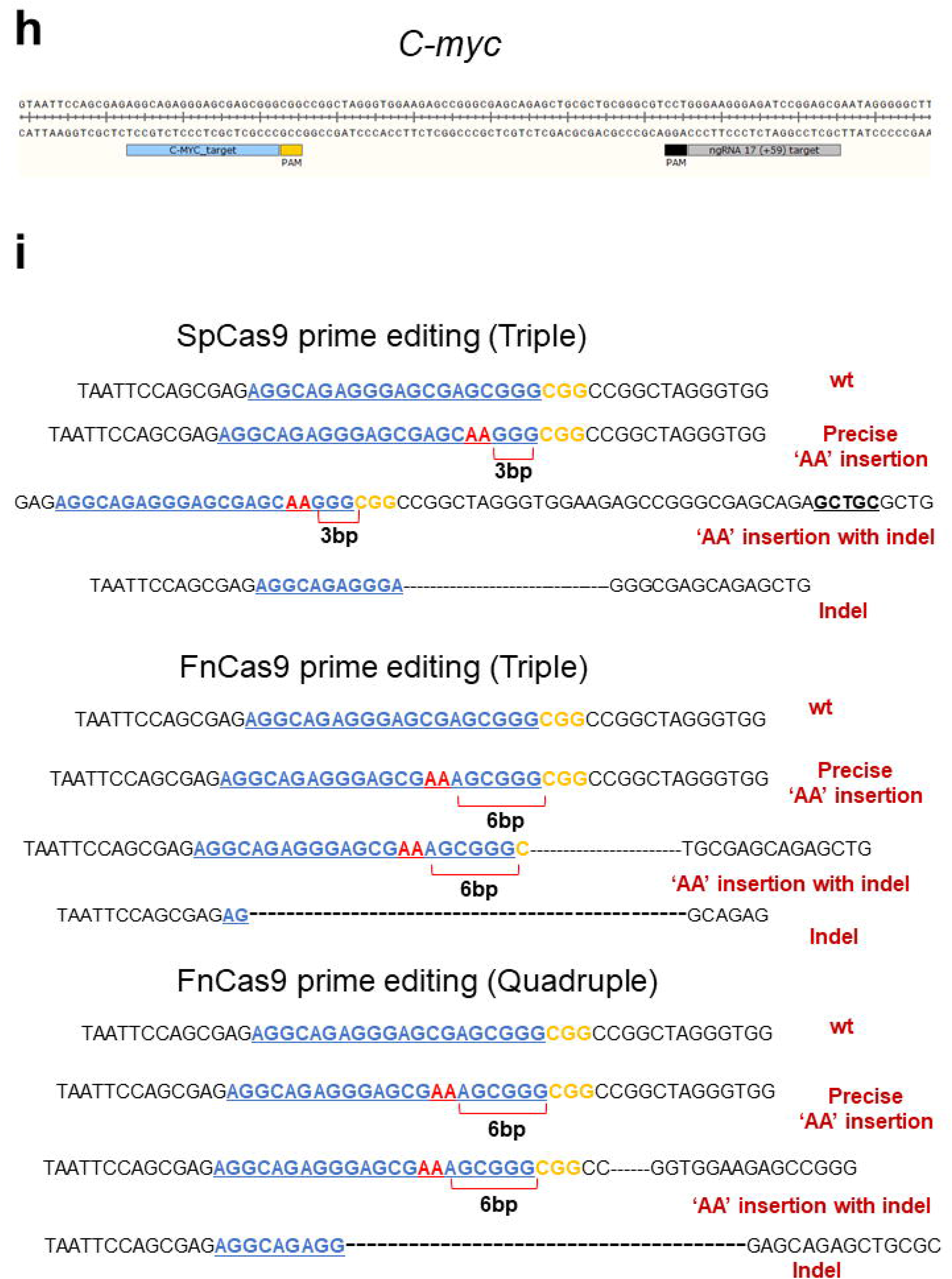
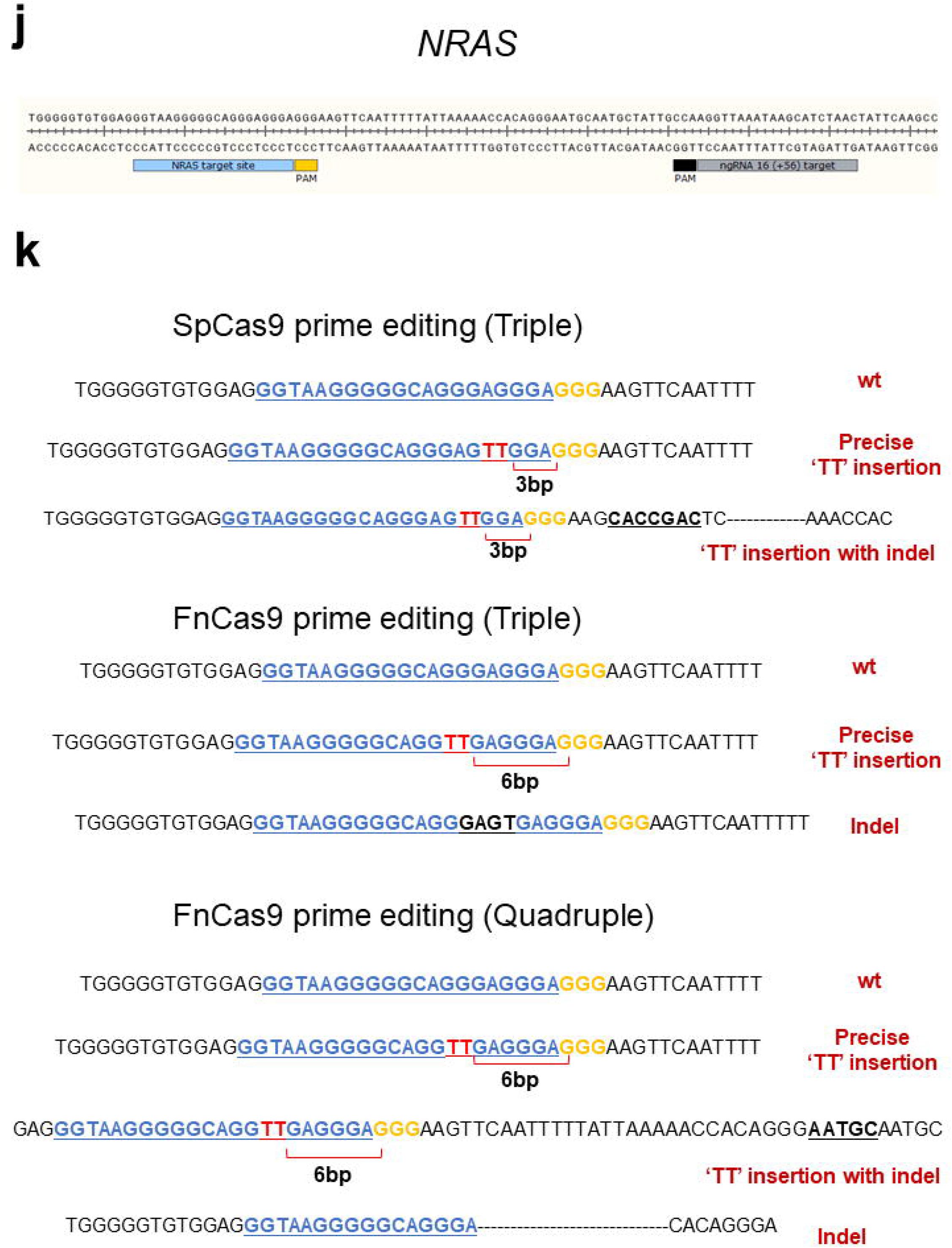

**Supplementary Figure 2.**
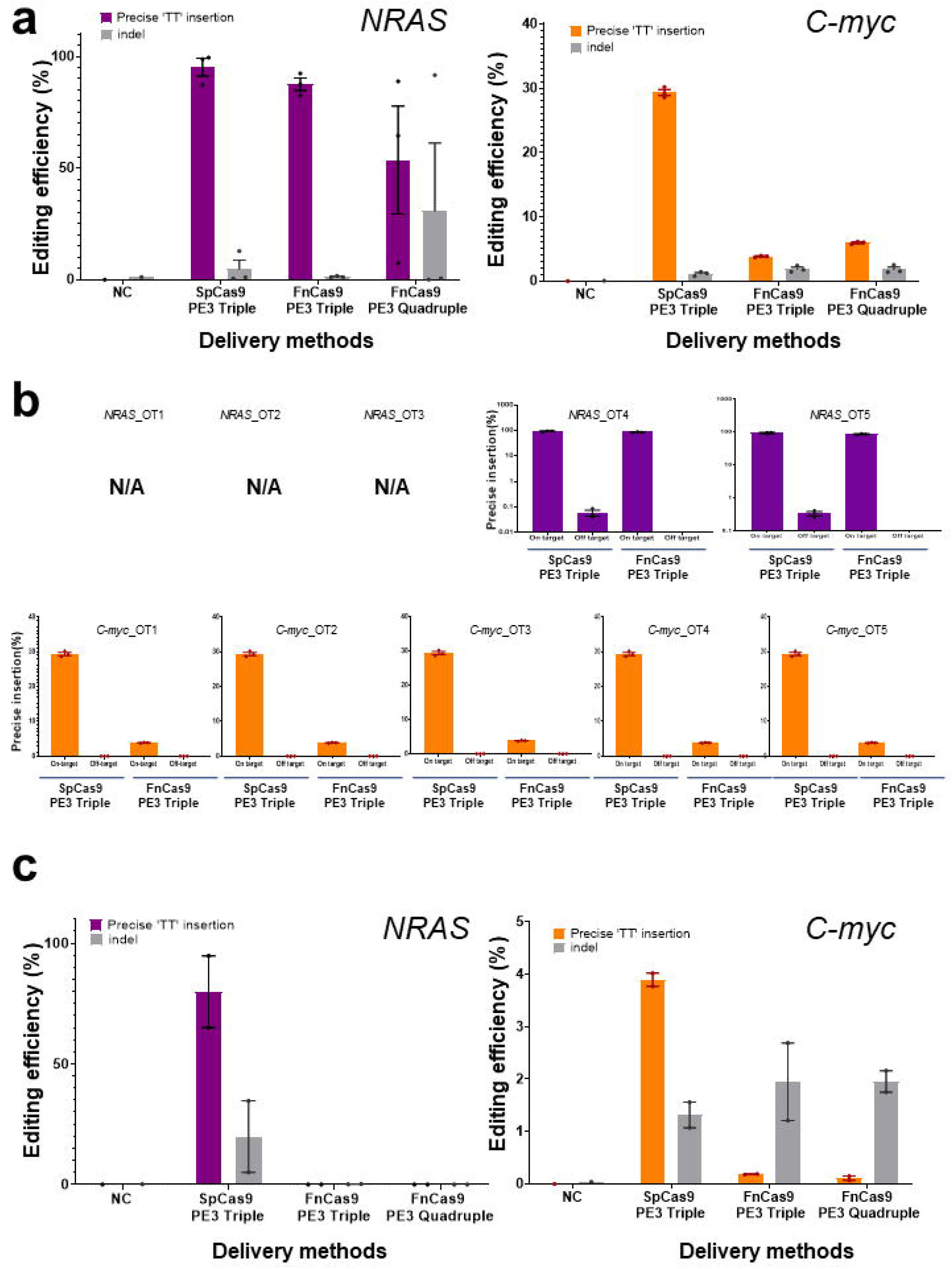
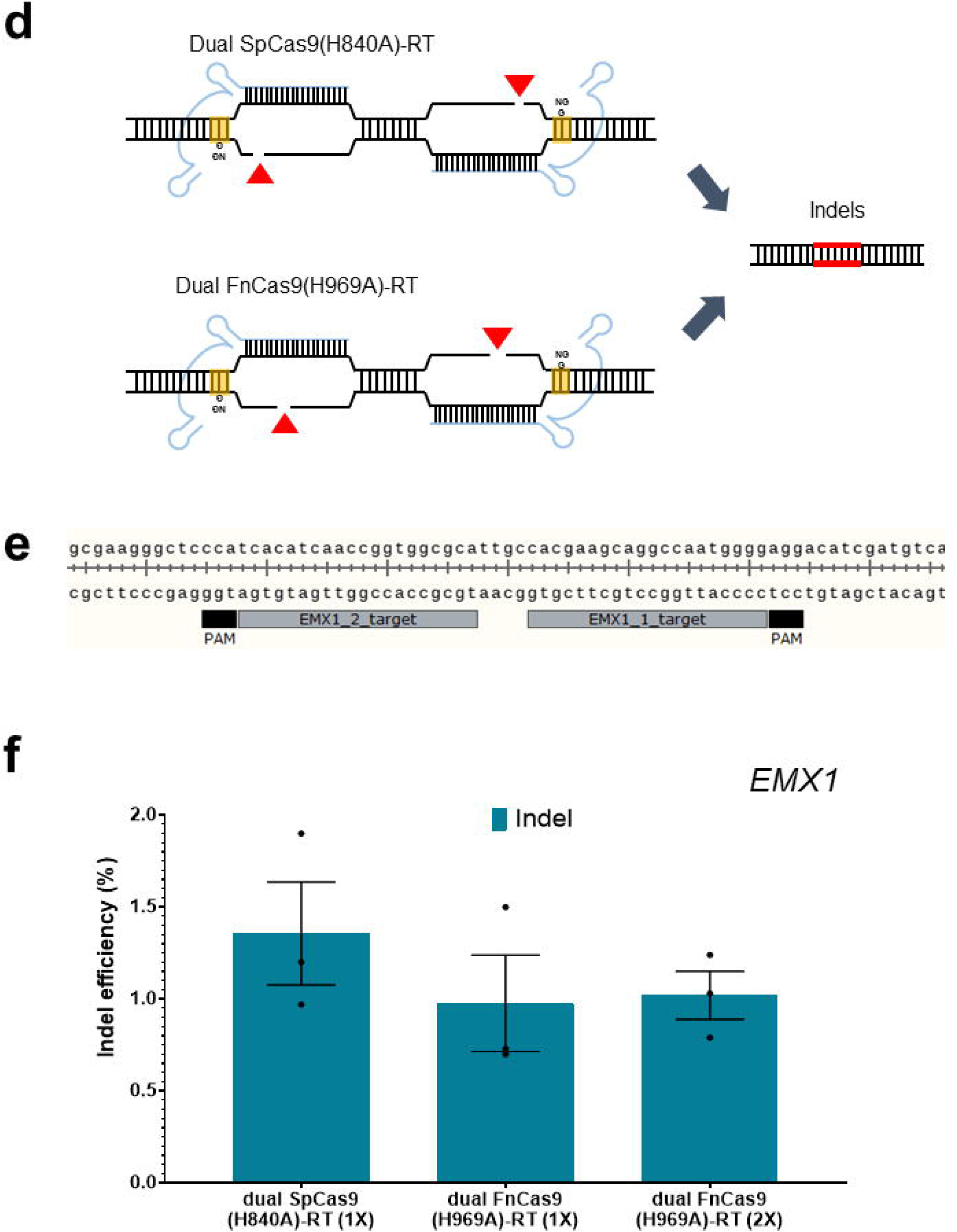
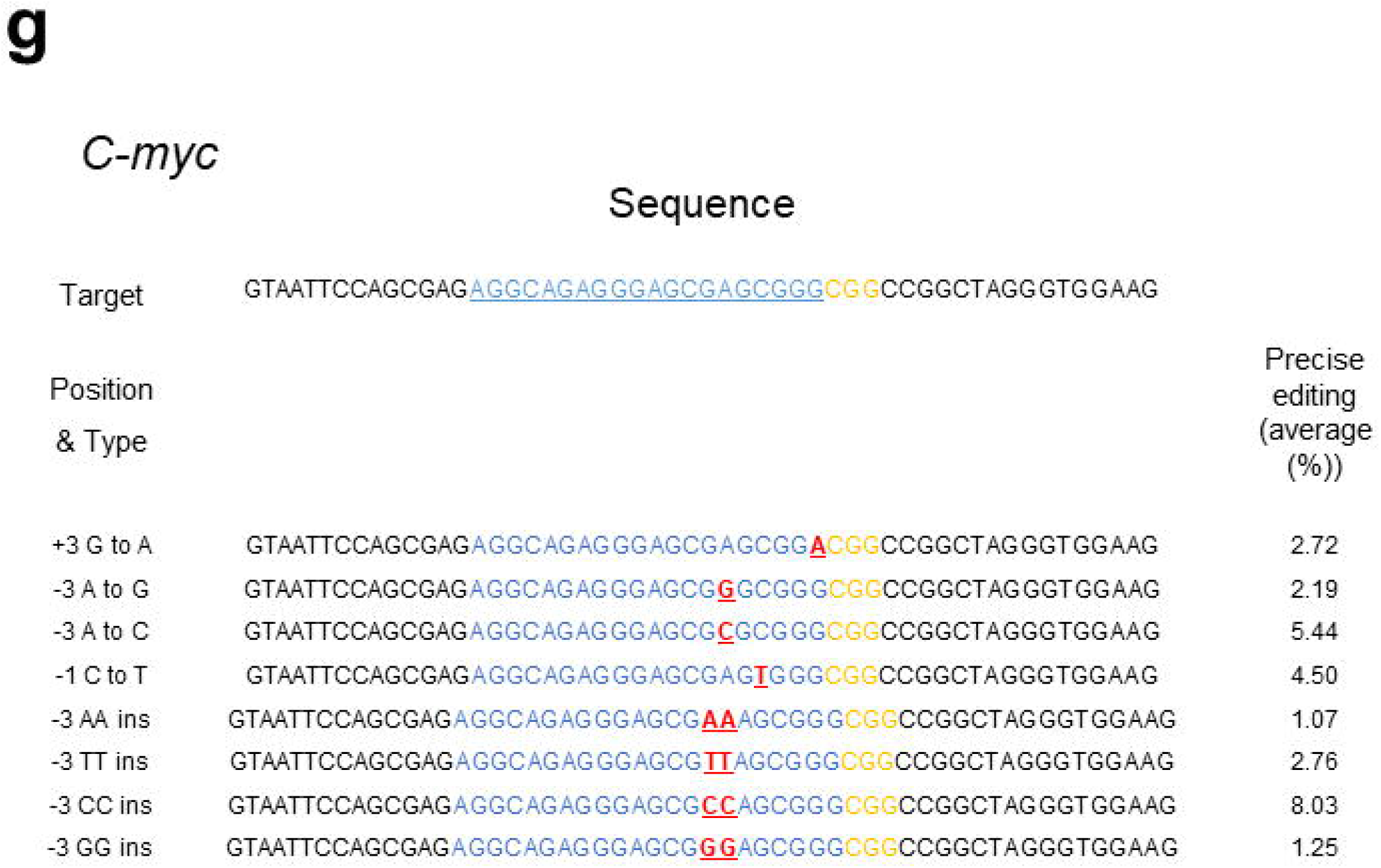

**Supplementary Figure 3.**
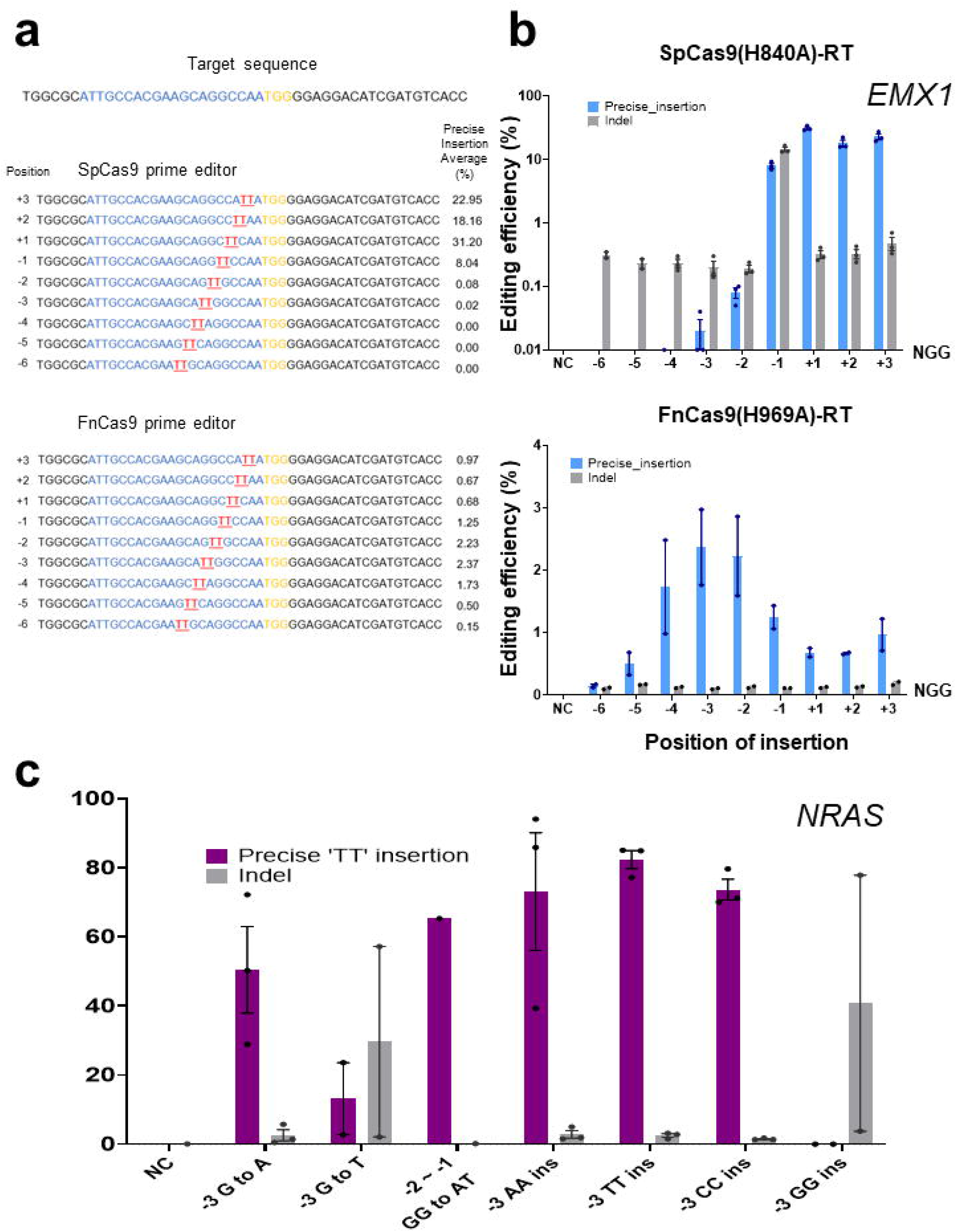
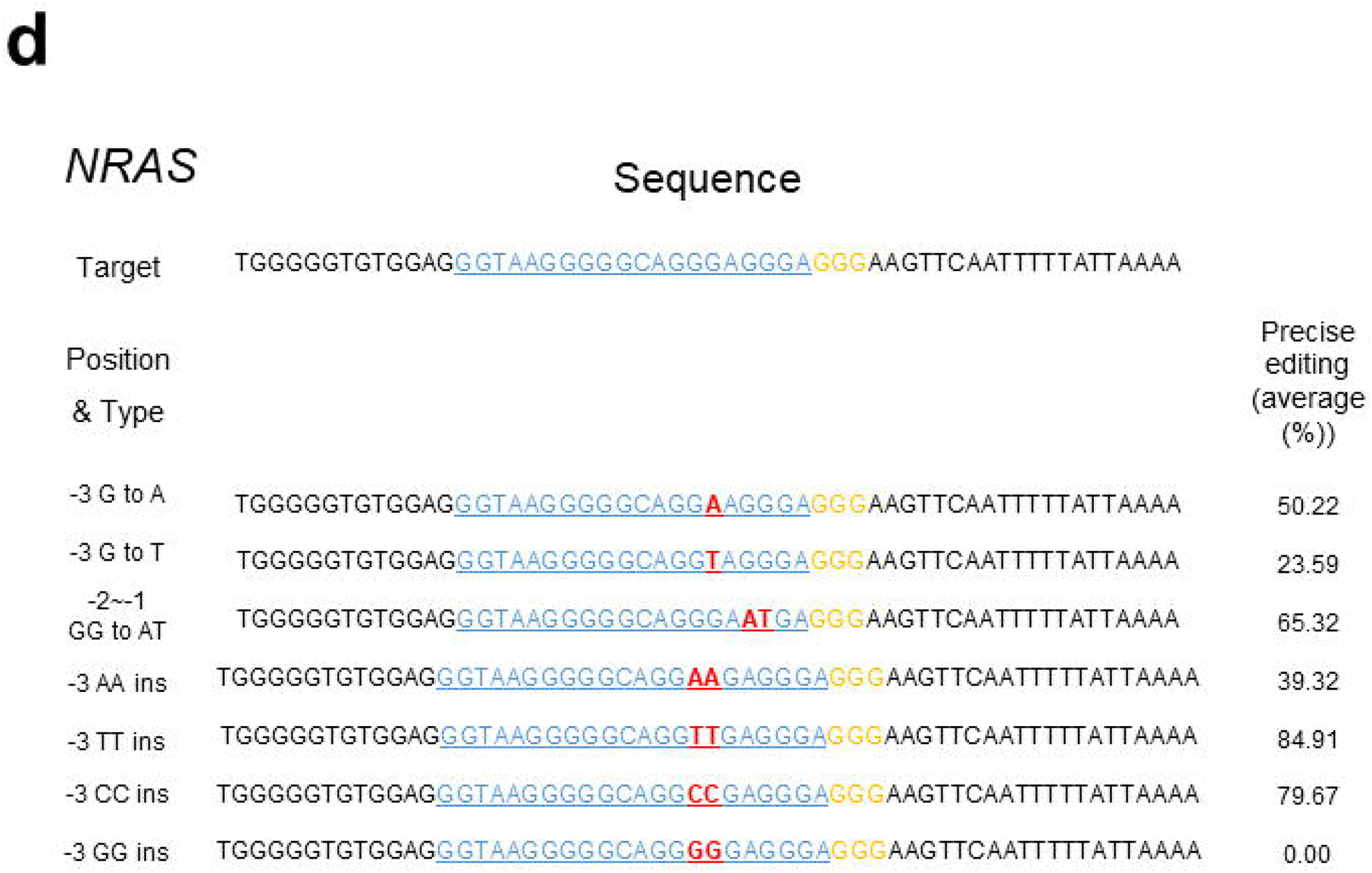

## Notes

### Competing Interest Statement

The authors have declared no competing interest.

